# Assessing exposure effects on gene expression

**DOI:** 10.1101/806554

**Authors:** Sarah A. Reifeis, Michael G. Hudgens, Mete Civelek, Karen L. Mohlke, Michael I. Love

**Affiliations:** Department of Biostatistics, University of North Carolina at Chapel Hill, Chapel Hill, NC, USA; Department of Biomedical Engineering, Center for Public Health Genomics, The University of Virginia, Charlottesville, VA, USA; Department of Genetics, University of North Carolina at Chapel Hill, Chapel Hill, NC, USA

**Keywords:** confounding, inverse probability weighting, observational genomics, parametric g-formula, regression

## Abstract

In observational genomics datasets, there is often confounding of the effect of an exposure on gene expression. To adjust for confounding when estimating the exposure effect, a common approach involves including potential confounders as covariates with the exposure in a regression model of gene expression. However, when the exposure and confounders interact to influence gene expression, the fitted regression model does not necessarily estimate the overall effect of the exposure. Using inverse probability weighting (IPW) or the parametric g-formula in these instances is straightforward to apply and yields consistent effect estimates. IPW can readily be integrated into a genomics data analysis pipeline with upstream data processing and normalization, while the g-formula can be implemented by making simple alterations to the regression model. The regression, IPW, and g-formula approaches to exposure effect estimation are compared herein using simulations; advantages and disadvantages of each approach are explored. The methods are applied to a case study estimating the effect of current smoking on gene expression in adipose tissue.

## Introduction

Increasing numbers of large-scale observational genomic datasets are available, in which tissue is collected from human donors sampled from a population. These donors differ in various ways, for example they may differ by age, sex, and other demographic variables, as well as by clinical or exposure variables such as body-mass index (BMI), diet, level of physical activity, history of medicine use, history of smoking or alcohol use, or various environmental exposures. Investigators are often interested in assessing the effect of various exposures on genomic variables, such as gene expression, as this may be useful to generate hypotheses about potential cellular mechanisms through which exposures may influence the development of diseases in human populations. Gene expression is a common molecular measurement in the context of exposure effects, although additional genomic assays, such as methylation, metabolites, protein abundance, may also be of interest.

A number of statistical methods have been proposed to address the problem of structural technical variation in gene expression measurements (Gagnon-Bartsch and Speed, 2012; Leek and Storey, 2007; Stegle et al., 2012), where *structural* refers to variation in the measurements across samples that is common across many genes. These methods address sample non-independence with a focus on estimation of latent factors, orthogonal to the biological condition of the samples, to be included in a linear model framework as regressors to account for the technical variation in the measurements. These methods can account for differences in measurements among sample preparation batches that may otherwise impair correct inference of differences across the biological conditions. Additionally, methods have been proposed to address sample correlations that arise from biological sources, for example repeated measures or genetically related individuals. Such sample non-independence can be addressed by explicit modeling of the known sample correlation structure as in a random effects framework (Cui et al., 2016); software for this approach include the *duplicateCorrelation* method (Smyth et al., 2005) in the *limma* package (Smyth, 2004), the *ShrinkBayes* package (Van De Wiel et al., 2012), or the *MACAU* package (Sun et al., 2017).

Regression frameworks alone may not be able to properly address the problem of confounding variables, whether measured or unmeasured, in observational datasets. Confounding is an issue when estimating causal effects, and as such is a distinct problem from the technical and biological sources of correlations among samples described above. Confounding has received relatively less attention in computational genomics, compared to the problems of structural technical variance or repeated measures. Existing work addressing confounding in observational genomic datasets has focused on sample matching (Heller et al., 2008), the combination of targeted minimum loss-based estimation (TMLE) and empirical Bayes shrinkage estimation (Hejazi et al., 2017), and TMLE for differential methylation controlling for observed methylation at neighboring genomic sites (Hejazi et al., 2018).

Since the exposure in an observational study is not randomly assigned, there is often confounding of the effect of exposure on the outcome. In general, randomized clinical trials to assess how various exposures affect gene expression cannot be conducted in human populations for ethical or feasibility reasons. Similar randomized studies can be performed on model organisms, but there is unique value in understanding the mechanism of these exposures in humans, and human populations are readily available for observational studies. In light of the increasing number of observational genomics studies on humans and the anticipated presence of confounding in such studies, methods of exposure effect estimation are worth further investigation.

For many studies, it is often useful to assess exposure effects on gene expression *in the exposed individuals*. This may be the case when researchers have particular interest in the effect of an exposure only on those types of individuals who likely will experience the exposure. For example, when studying the effect of smoking, it is often most relevant to obtain effect estimates interpretable for those who actually smoke, as opposed to the effect smoking would have, averaging over all the persons in the general population. In these cases, the target estimand is referenced as the exposure effect in the exposed; this terminology will be used interchangeably with the average treatment effect in the treated (ATT) throughout. In contrast, the average treatment effect (ATE) is a different estimand and is interpretable in the context of the population as a whole, including both treated and untreated individuals. This paper will focus on obtaining estimates for the former, the ATT.

The conventional approach to quantifying the effect of exposure, while attempting to adjust for confounding, is to fit a linear model of gene expression with the exposure and various potential confounders as covariates. In this approach, the estimated coefficient of the exposure variable is often interpreted as an estimate of the exposure effect. However, fitting the conventional linear model falls short of our goal in two respects: (1) it does not directly produce an estimate of the effect of interest, the effect of the exposure *in the exposed individuals*, who may differ in various respects from unexposed individuals, and (2) it may not appropriately adjust for confounding, resulting in estimates that are not consistent and confidence intervals that do not provide their nominal coverage. For these reasons, this paper demonstrates existing causal inference methods that can be employed in these scenarios to adequately adjust for confounding and return consistent exposure effect estimates and valid confidence intervals.

Regression, inverse probability weighting (IPW), and the parametric g-formula are compared herein for obtaining exposure effect estimates. Both IPW and the parametric g-formula are methods established and widely applied to observational studies in the causal inference and epidemiology literature. This paper seeks to demonstrate that these methods also have utility in the space of observational genomics. In general, IPW uses weights to construct a “pseudo-population” in which there is no longer confounding of the effect of an exposure on the outcome of interest; simple linear regression is then applied to this “pseudo-population” to obtain consistent estimates of the exposure effect (Robins et al., 2000). The parametric g-formula, also referred to as standardization, entails fitting an outcome regression model and then averaging the predicted outcomes across all individuals for a fixed level of the exposure. Both IPW and the parametric g-formula rely on standard assumptions of causal inference (conditional exchangeability, positivity, and consistency), but they differ in the modeling assumptions required (Naimi et al., 2017). Statistical validity of the IPW and parametric g-formula methods rely on asymptotic justifications, and are not guaranteed to perform well for small sample sizes.

The following is an outline for the remaining sections of this paper. A brief summary is given of the models used, followed by more extensive description in the Methods section for both the simulation study and data analysis; formal definitions are left to the Supplementary Methods. In the Results section, the methods are compared in simulations and a data analysis. Simulation studies are based on the Metabolic Syndrome in Men Finnish cohort (METSIM) (Laakso et al., 2017) analysis dataset (n = 770). In particular, scenarios falling into three categories for a binary exposure effect on gene expression are investigated: no confounding, and confounding both with and without interaction(s) between exposure and covariates. The METSIM cohort data is then analyzed to investigate the effect of current smoking on gene expression in adipose tissue. The Discussion section concludes with reviewing and providing insight into the main results, and addresses limitations of and future directions for this research. The Supplementary Methods gives details on the models for the regression, IPW, and g-formula approaches, as well as estimates of standard errors and assumptions required for each approach. In the Supplementary Results, it is first established why the proposed methods are being compared to linear regression alone, and not to the linear model with empirical Bayes moderation of the standard errors (for example, as implemented in the *limma* package (Smyth, 2004)). The Supplementary Results section concludes with sensitivity analyses for the METSIM cohort data. Appendix A demonstrates the equivalence of the g-formula estimator presented in the main text and the g-computation algorithm of Snowden et al. (2010). Appendix B provides R markdown (Rmd) workflows showing the generation of the simulated data and performing the three methods on a simulated dataset.

## Methods

### Summary of models compared

Three exposure effect estimation approaches were assessed in the following evaluation: traditional linear regression, inverse probability weighting, and the parametric g-formula. In fitting the models associated with each approach, it was assumed that the models were correctly specified, the set of confounders identified was sufficient to adjust for confounding, and the data were free from selection bias and systematic measurement error. The regression model with the exposure and potential confounders as predictors was fit using ordinary least squares for both the parameter estimates and their standard errors, in keeping with the conventional approach. For the IPW approach, the confounders were used as predictors in a logistic regression model of the exposure to obtain the weights, and the weights were used in the simple linear regression model of gene expression on the exposure using weighted least squares. In the g-formula approach, all potential confounders were centered at the mean value in the exposed, and the linear regression model with the exposure, centered confounders, and their interactions was fit using ordinary least squares. The standard errors for both the IPW and g-formula estimators were computed using stacked estimating equations (Stefanski and Boos, 2002). The details of using these methods with observational genomics data are described further in the Methods Section, formally defined in the Supplementary Methods section that follows the main body of the paper, and demonstrated in the R code included in Appendix B.

### Simulation Study

Performance of methods was first compared using a simulation. The simulated covariates and exposure were based on counterparts from the METSIM data analysis in the next section. Specifically, the simulated variables included a current smoking indicator and five variables considered to be potential confounders of the relationship between current smoking and gene expression. Table 1 below gives more details regarding variable distributions and dependencies.

**Table 1:**
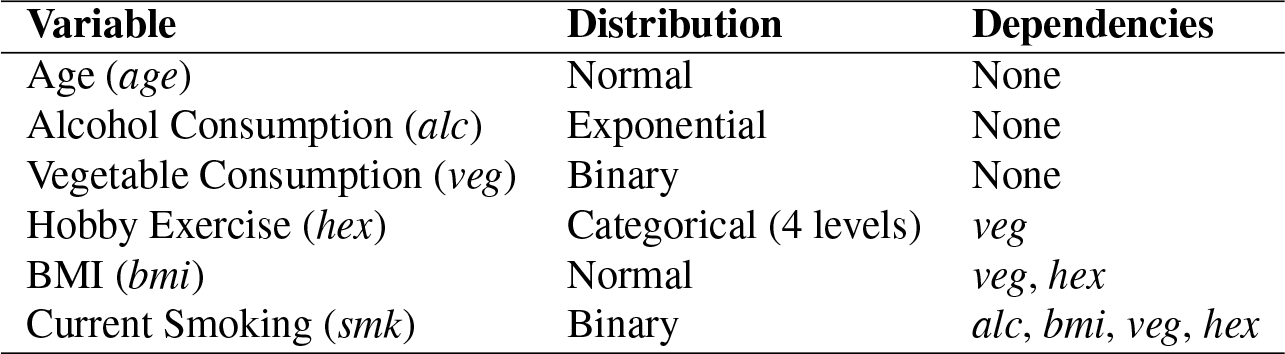
Definitions of exposure and confounding variables for simulation study comparing regression, IPW, and the parametric g-formula

Following the generation of the variables in Table 1, normalized gene expression values for various scenarios were simulated as well. For scenarios where no confounding was present, the mean of the expression values were dependent on only the exposure or none of the variables. When confounding was present, the mean expression values were dependent on both the exposure and the other covariates, with some scenarios including interactions between the exposure and the covariates. Expression values were generated with different means for each individual and each gene, according to the simulated exposure and covariates. The mean for gene *g* and individual *i* was

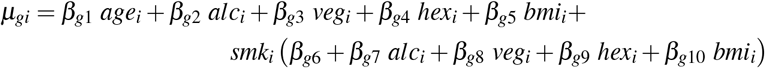

where the *β*_*gk*_, *k* = 1, …, 10, varied by gene and were restricted to *β*_*gk*_ ∈ [−2, 2] for this simulation study. Here *age*, *alc*, and *bmi* were each centered about their population mean and scaled. Note that there was no quadratic age term and no interaction term for age and smoking in the true model for mean expression, although the former was included in the analysis models; these terms can be thought of as not contributing to the true mean gene expression for any individual. The standard deviation of each gene was set to the same value to aid with comparing results for different genes, and was equal to 0.24.

The variables listed in Table 1 were generated for a population of 10 million individuals, from which the true ATT was calculated for each gene. From this population, 1000 sets of 770 individuals were randomly selected with replacement to build the analysis datasets. The sampling algorithm allowed for the same individual to be present in more than one dataset, but not more than once within a single dataset.

In all instances the regression model had the same form shown in equation S.1 of the supplementary materials, namely the exposure and covariates (with both linear and quadratic age terms) were each present as main effects in the model and no interaction terms were included; the covariates *age*, *age*^2^, *alc* and *bmi* were centered at their sample mean and scaled for each dataset, as would typically be done to avoid collinearity with the intercept. The standard errors used to construct the 95% confidence intervals were obtained through fitting the model with ordinary least squares.

For the IPW approach, first the logistic regression model in equation S.2 of the supplementary materials was used to compute the components needed for the weights for each dataset, with terms for the five covariates and a quadratic age term. Weights for the ATT were then constructed according to expression S.3 of the supplementary materials. Then the linear regression model of gene expression in S.4, with only the exposure and an intercept, was fit with the weights to obtain the effect estimate. Standard error estimates used to obtain the 95% confidence intervals were computed with the stacked estimating equations approach using the *geex* package (Saul and Hudgens, 2017) in R, taking into account estimation of the weights by stacking the estimating equations for the logistic regression model with those used in computing the IPW estimator.

The regression model given in equation S.5 of the supplementary materials was used to obtain the g-formula estimate for each dataset. It contained variables for the main effects of the exposure and covariates (including both linear and quadratic age) as well as interactions between *smk* and each of *bmi, veg, hex*, and *alc*. Since the ATT is being compared across methods in this simulation study, the non-exposure covariates were all centered at the sample mean in the exposed. The standard error estimates used for the 95% confidence intervals were computed with stacked estimating equations by passing *geex* the set of stacked estimating equations corresponding to the covariate means and the regression model parameters.

### METSIM Smoking Exposure Effect

In the METSIM project dataset, the goal was to obtain the estimated effect of current smoking on gene expression in the smokers, adjusting for the set of potential confounders: linear and quadratic age, BMI, alcohol consumption, vegetable consumption, and hobby exercise. The data consisted of adipose gene expression values and several covariates, measured for *n* = 770 individuals (details on data preprocessing in Supplementary Methods). There were no missing outcome, exposure, or covariate values. This cohort was analyzed in Civelek et al. (2017), where BMI and linear and quadratic age were considered confounding variables for various phenotypic traits. This analysis, and consultations with a subject-matter expert, guided the choice of the confounding variable set. Note that current smoking was not examined by Civelek et al., so their analyses were not compared directly to those in this paper. Each of the three methods introduced above were implemented for this cohort in the analyses that follow.

Table 2 briefly summarizes the variables used in this analysis; the range, mean and standard deviation are reported for continuous variables and the levels and distribution are given for each categorical variable. For hobby exercise, higher levels denote increased activity levels.

**Table 2:** Descriptive statistics for the METSIM cohort data.

| Variable | Range | Mean (SD) |
| --- | --- | --- |
| Age (*yrs*) | (45, 68) | 55 (5) |
| Alcohol Consumption (*g/wk*) | (0, 1134) | 105 (119) |
| BMI (*kg/m<sup>2</sup>*) | (18.5, 48.1) | 26.6 (3.5) |
|  | Level | Proportion |
| Vegetable Consumption | Everyday | 0.83 |
|  | Not Everyday | 0.17 |
| Hobby Exercise | 1 | 0.05 |
|  | 2 | 0.28 |
|  | 3 | 0.18 |
|  | 4 | 0.49 |
| Current Smoking | Yes | 0.17 |
|  | No | 0.83 |

Note that the models used for the data analysis had the same form as those fit for the simulation study, described in the section above. The estimated exposure effect was obtained using regression, IPW, and the g-formula for each of the 18,510 genes; the coefficient for current smoking represented the exposure effect in each model. The models were fit and standard errors computed, again using the same process as for the simulation study. With each approach, a *t*-test of no effect of exposure on gene expression was performed for each gene and the resulting p-values were adjusted using the correction from Benjamini and Hochberg (1995) to control the false discovery rate.

To compute weights for the 770 individuals, the logistic regression model of current smoking was fit with the previously listed covariates as predictors. Before computing effect estimates and standard errors, it is good practice to check that the weights have a mean value close to the expected value (details in Supplementary Methods) and that none of the weights are extreme. The weights had mean value 0.34, which was exactly what was expected for these data. There was one weight with a value of 5.26, substantially larger than the rest; for this reason, a sensitivity analysis was conducted in the Supplementary Results section where the observation with this large weight was deleted and the same analysis was performed again to investigate the influence of this observation.

For both the g-formula and regression methods, leverage values were computed for each observation to determine if there were any influential points in these analyses; the same observation returned the largest leverage point for both the g-formula and regression, which took values 0.38 and 0.11 respectively. In both instances these leverage values were approximately twice the magnitude of the next largest value, so another sensitivity analysis was conducted in the Supplementary Results where the observation with this large leverage value was deleted and the analysis was performed again to investigate its influence.

## Results

### Simulation Study

The empirical bias and confidence interval coverage and width for the regression, IPW, and g-formula estimators are shown in Table 3, averaging over 1000 simulations per scenario. While there were instances where all estimators appeared to perform well, the IPW and g-formula estimators provided advantages over regression in a subset of the simulated scenarios. In particular, the IPW and g-formula estimators remain unbiased and meet nominal confidence interval coverage in all scenarios, but the regression estimator manifests bias and fails to meet nominal coverage in the presence of exposure-covariate interactions. The reported coverage represents the proportion of confidence intervals which included the true value of the ATT. The true ATT value is shown for each scenario, and is based on the original population of 10 million individuals. The null case where there was no effect of current smoking on gene expression is shown in addition to several representative scenarios where confounding was present, two without and three with one or more interactions between exposure and confounders.

**Table 3:** Average empirical bias and 95% confidence interval (CI) coverage and width for the regression (Reg), IPW, and g-formula (G-form) estimators.

| Scenario | True | Estimate Bias |  |  | 95% CI Coverage |  |  | 95% CI Width |  |  |
| --- | --- | --- | --- | --- | --- | --- | --- | --- | --- | --- |
|  | ATT | Reg | IPW | G-form | Reg | IPW | G-form | Reg | IPW | G-form |
| Null Case | 0.00 | 0.00 | 0.00 | 0.00 | 0.96 | 0.96 | 0.95 | 0.10 | 0.10 | 0.10 |
| No Interactions 1 | -2.00 | -0.00 | 0.00 | 0.00 | 0.95 | 0.96 | 0.95 | 0.10 | 0.11 | 0.10 |
| No Interactions 2 | 2.00 | 0.02 | 0.01 | -0.00 | 0.91 | 0.92 | 0.94 | 0.15 | 0.35 | 0.18 |
| Interactions 1 | 1.59 | 0.12 | 0.00 | 0.00 | 0.34 | 0.96 | 0.96 | 0.18 | 0.39 | 0.39 |
| Interactions 2 | -0.36 | -0.20 | 0.00 | 0.00 | 0.19 | 0.96 | 0.95 | 0.23 | 0.51 | 0.51 |
| Interactions 3 | -1.75 | -0.20 | 0.01 | 0.01 | 0.37 | 0.95 | 0.95 | 0.31 | 0.70 | 0.70 |

True ATT values in these simulations are in units of log_2_ fold change in gene expression. The most extreme scenario shown here, where the true ATT is 2.00, thus represents a 4-fold change in gene expression attributable to smoking. Additional simulation studies were conducted that are not shown in this table, but the results were similar to those included. Across all simulations run, the average bias of the regression estimator took values in the range [−0.29, 0.29], whereas the IPW and g-formula average biases were contained to [−0.01, 0.01]. For this simulation study setup, the regression bias appears to be larger in magnitude when interaction terms involving alcohol and hobby exercise contributed to the true ATT.

When smoking and the confounders did not interact to influence gene expression, e.g., the first three examples in Table 3, all methods met nominal coverage and yielded no bias on average. In scenarios where interactions existed between smoking and the confounders, the regression effect estimator had nonzero average bias and substantially below nominal confidence interval coverage. The IPW and g-formula estimators both resulted in very low or no bias on average, and both uniformly met (or very nearly met) the nominal confidence interval coverage. Except in the null case, the IPW and g-formula estimators were generally more variable than the regression estimator. When smoking and the confounders interacted, the IPW and g-formula estimates had confidence intervals that were on average approximately twice as wide as those for regression. Of the two methods that overall maintained the nominal coverage across scenarios, IPW and g-formula, they tended to have comparable interval width except in one of the no-interaction scenarios, in which IPW had nearly double the average interval width of g-formula.

### METSIM Smoking Exposure Effect

Regression, IPW, and g-formula were applied in order to assess the effect of smoking on gene expression among smokers in the METSIM cohort. Estimates and confidence intervals for each of the three methods, for the top two genes as ranked by p-value are presented in Figure 1a - 1b (the top two genes are shown as their effect sizes and test statistics were appreciably larger compared to the other genes). The three methods were in agreement on the ranking for the top three genes, but beyond this the rankings were not consistent across method (Figure 1c). The top two genes in terms of estimated effect size and test statistic were *CYP1A1* and *CYP1B1*, which were expected as they play a role in metabolizing cigarette smoke (Nebert and Russell, 2002). The confidence intervals for the IPW and g-formula estimates of the top two genes were similar in width, and the regression confidence interval was substantially less wide. While the −log_10_ adjusted p-values for the top two genes were all large for each of the three methods (highly significant), those from regression were much larger than those from IPW and the g-formula (*CYP1A1*: R=119, W=44, G=44; *CYP1B1*: R=30, W=18, G=20). This is in accordance with the displayed difference in confidence interval widths for the smoking effect estimates of these two genes. Although the ATT estimates were similar across methods for the genes shown, instances of substantial differences in standard error produced vastly different confidence intervals and p-values.

**Figure 1:**
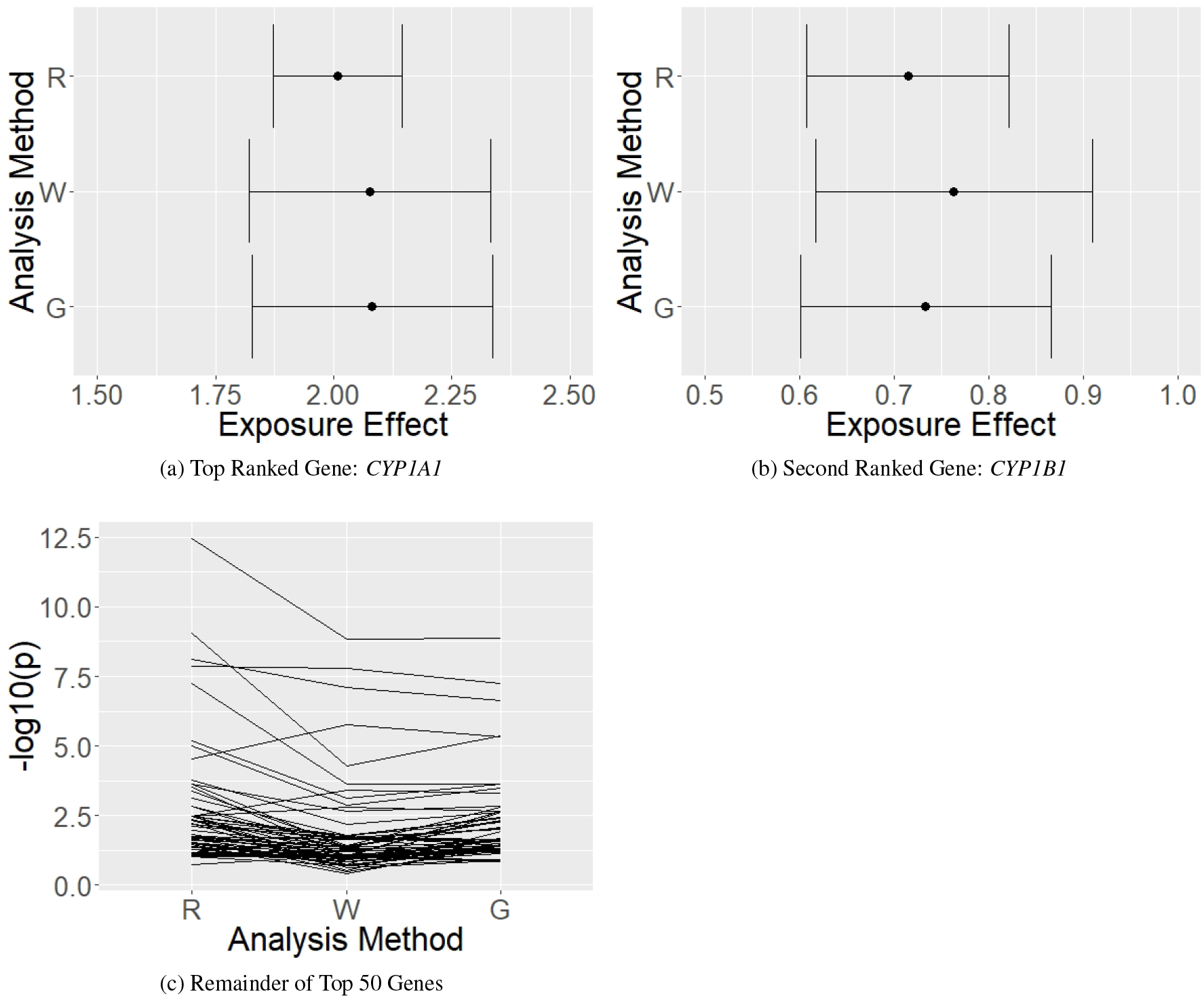
METSIM analysis results for the top 50 genes, ranked by p-value. (a)-(b) Estimates and 95% confidence intervals for the effect of current smoking in the smokers for genes *CYP1A1* and *CYP1B1*, respectively. Note that the null value for the smoking effect estimate (log_2_ fold change = 0) is not included on the x-axis. (c) −log_10_ adjusted p-values for the top 50 genes (omitting top 2), for each of R = Regression, W = Inverse Probability Weighting, and G = Parametric G-Formula.

IPW and the g-formula tended to produce larger p-values than regression for these data, though this was not the case for every gene (Figure 1c). Note that in this figure the top two genes were not represented, as their −log_10_ adjusted p-values were much larger than the others. Additionally, the smoking effect estimates for all genes except the top two were in the range (−0.5, 0.5), with many very close to zero.

## Discussion

Geneticists and epidemiologists may analyze differential gene expression due to exposures in a population in order to generate hypotheses as to how those exposures may be related to health outcomes. Results of these analyses, i.e., lists of genes affected by the exposure under a false discovery rate bound and their associated effect sizes, may be inaccurate and irreproducible across study populations unless potential confounders of the exposure and gene expression are properly adjusted for. Here, exposure effect estimates were compared using causal inference techniques such as IPW and the parametric g-formula, as well as with common practice techniques such as regression. Comparisons were performed across simulated data and a data analysis in which gene expression was measured in subcutaneous adipose tissue. Tissue donors also had various clinical and demographic covariates measured, and it was desired to adjust for differences among the exposed and unexposed donors when estimating the ATT. Analyses of the METSIM cohort found that estimation method did not make a substantial difference for the effect estimates for the top two genes, *CYP1A1* and *CYP1B1*. Simulations based on the METSIM data showed that there was potential for the regression estimate to be biased, but the effect biases observed in the data analyses were small. Differences between the methods were most pronounced when examining the standard errors and therefore also the resulting confidence intervals and p-values. In particular, simulations based on the METSIM data showed that if there were any interactions of the exposure with the confounder(s), the regression method produced confidence intervals that can have far below nominal coverage. Furthermore, what may appear to be modest changes in standard errors can produce a dramatically different set of adjusted p-values for a given set of genes.

In addition to the standard errors, regression as applied here differs from IPW and the parametric g-formula in that the regression estimates do not represent the effect of exposure in the exposed, but rather in the population as a whole. While IPW and the g-formula can be adapted to produce the exposure effect in the population or sub-population of interest, regression estimates remain population-wide estimates — unless operating under the assumption that ATE and ATT are equal.

It should be mentioned that if all appropriate interaction terms were included in the regression model, the parameter estimates could be combined to yield consistent, conditional exposure effect estimates. However, this approach was not taken here for two reasons: (i) the goal was to obtain one exposure effect estimate that could be read directly from software output without additional steps, and (ii) the exposure effect estimate constructed via combination of exposure and interaction terms would be interpreted conditional on values of the covariates, whereas the desired exposure effect estimate has a marginal interpretation. Expanding on this second reason, regression with all appropriate interaction terms would result in a variety of exposure effects across combinations of covariates used for conditioning, as opposed to the causal approaches presented here which provide one exposure effect integrating over the exposed individuals. If all appropriate interaction terms were included in the regression model and centered at the mean in the population of interest, then the regression model would be equivalent to the parametric g-formula model.

Often in analyses of exposure effects on microarray gene expression, the *limma* method is used to fit the regression models and obtain a moderated *t*-statistic (Smyth, 2004), whereas here the ordinary *t*-statistic was used. The simulation results illustrating the rationale behind this choice are included in the Supplementary Results section. In short, the sample size of the cohort analyzed here (*n* = 770) was sufficiently large that the ordinary and moderated *t*-statistics are practically equivalent. More recently, another method has been proposed involving the combination of TMLE and empirical Bayes shrinkage estimation, which has demonstrated utility with small and moderate sample sizes (Hejazi et al., 2017). The intended audience of this paper is working with larger datasets, allowing for reliance on large sample theory; for this reason the simpler and more readily available approach was used for the regression analysis. Here, pre-normalized microarray gene expression, which takes continuous values, was analyzed, while RNA sequencing experiments result in count-valued observations for gene expression. In order for the causal inference approaches shown here to be applied to count data from RNA sequencing, it would be desirable to first perform library size scaling and apply a variance stabilizing transformation to the gene expression (Anders and Huber, 2010; Law et al., 2014), though such datasets and procedures were not evaluated in the present work.

The IPW and parametric g-formula approaches are both presented here as alternates to regression that adequately adjust for confounding in a wider variety of circumstances. While IPW and the g-formula both accomplish this goal, they require slightly different assumptions and they have different strengths and weaknesses. The IPW estimator relies on correct specification of the exposure model, which is often more plausible than correct specification of the outcome model. IPW can be sensitive to extreme weights, as shown in the sensitivity analysis results for the METSIM data, and can be more variable than the g-formula estimator. While the consistency of the g-formula estimator relies heavily on the correct specification of the outcome model, it appears to be less sensitive to extreme values of the covariates and can be less variable than the IPW estimator. Due to the limited overall differences in the bias and efficiency of the IPW and g-formula estimators, the researcher is encouraged to choose among methods based on their relative confidence in specification of the exposure or outcome models.

There are several assumptions made in these analyses which may be violated and deserve further exploration. Firstly, the assumption of causal consistency states that there are not multiple ways to be a current smoker. This assumption is clearly not met since the amount of cigarettes smoked daily can vary from person to person, but this assumption can be replaced by another less stringent assumption. In particular, it can more reasonably be assumed that these different versions of exposure do not have any bearing on the causal effect; this is referred to as treatment variation irrelevance. The data analyses above rely on the additional assumptions that the set of confounders *L* are sufficient to adjust for confounding, and that there is no systematic measurement error or selection bias influencing these data. If any of these assumptions are unmet then the exposure effect estimates may be biased. Furthermore, formal arguments for the methods presented rely on large sample theory; while there is some empirical evidence suggesting that, for example, IPW can perform well with moderate sample sizes (Pirracchio et al., 2012), these methods are not guaranteed to perform well for small or moderate samples.

The performance of doubly robust estimators for estimating exposure effects on gene expression could be investigated in future work. Doubly robust estimators have been shown to provide advantages over IPW or the g-formula (Lunceford and Davidian, 2004; Moodie et al., 2018; Naimi and Kennedy, 2017), and could conceivably allow a relaxation of certain modeling assumptions in observational genomics analyses while maintaining the desirable properties of causal methods. Additionally, this paper focuses on binary exposures but future work could expand this to allow for continuous or longitudinal exposures.

## Acknowledgments

The authors thank the following individuals for useful discussions: Nathan Kallus, Melissa Troester, the METSIM investigators. S.A.R. was supported by The Chancellor’s Fellowship from The Graduate School at the University of North Carolina at Chapel Hill. M.G.H. was supported by R01 AI085073. M.C. was supported by R01 DK118287 and R21 HL135230. K.L.M. was supported by R01 DK093757. M.I.L. was supported by R01 HG009937, R01 MH118349, and P30 ES010126.

## Data Availability

The data that support the findings of this study are openly available in GEO under accession number GSE70353 (https://www.ncbi.nlm.nih.gov/geo/query/acc.cgi?acc=GSE70353) (Civelek et al., 2017).

## Appendix A Equivalence of g-formula estimator and Snowden’s g-computation algorithm

Proof that the g-formula estimation method presented in this paper is equivalent, in this setting, to the g-computation algorithm outlined by Snowden et al. (2010) for both the ATE and the ATT.

Let *Y* be an outcome variable and *A* be a binary treatment variable. Assume *L* is a vector of length *J* which represents a sufficient set for confounding adjustment. Consider the following linear model

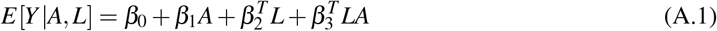

where *β*_2_*, β*_3_ are parameter vectors of length *J*. Model A.1 is assumed to be correctly specified. The parameter estimates 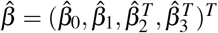 are found by solving

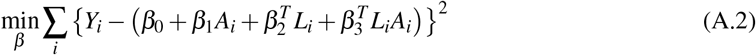

where 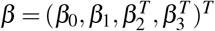. The Snowden g-formula estimator for the ATE is obtained using the parameter estimates 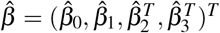 to compute the predicted values under the counterfactual scenarios of no treatment (*a* = 0) and treatment (*a* = 1) for all. Specifically, let 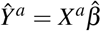 denote the vector of predicted values for each individual under the counterfactual scenario that all individuals receive treatment *a*, and where the design matrix *X*^*a*^ has rows of the form 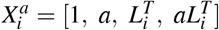 for *i* = 1, …, *n* and *a* = 0, 1. The Snowden g-formula estimator of the ATE can be expressed

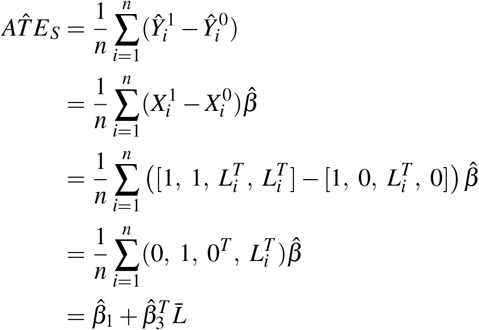

where 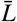 is a vector of length *J* with elements equal to the sample means of the *J* confounding variables.

Now, let 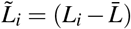 and 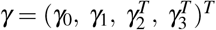. Consider finding 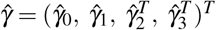 which solves

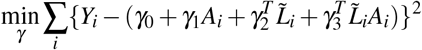

or equivalently

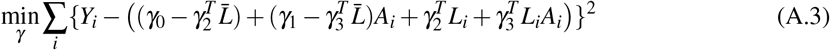

We can find 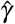 by first reparameterizing. Let 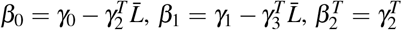, and 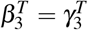. This optimization problem is then equivalent to solving A.2 above, which yields the usual least squares estimator 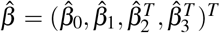. Therefore, 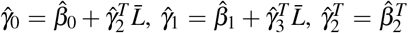, and 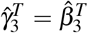. That is, 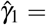 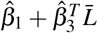. Note that 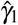 is the estimated exposure coefficient from the g-formula model proposed in the Supplementary Methods for the ATE, and the right side of this equality is exactly 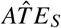 from above. The equivalence of the two g-formula estimators has thus been shown for the ATE.

The equivalence proof for the ATT estimators is analogous to the ATE proof above. Wang et al. (2017) propose an extension of Snowden et al.’s g-computation algorithm for the ATT; they show that by restricting *X*^*a*^ to only the treated individuals (i.e., those with *A* = 1), the algorithm returns a consistent estimate of the ATT. In particular, let 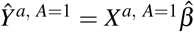 denote the vector of predicted values for the *n*_1_ individuals with *A* = 1 under the counterfactual scenario that these individuals receive treatment *a*, and where the design matrix *X*^*a, A*=1^ has *n*_1_ rows with the same form given above. Here 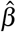 remains the usual least squares estimator found by solving A.2 above. The Wang et al. (2017) extension to the Snowden g-formula estimator for the ATT can be expressed

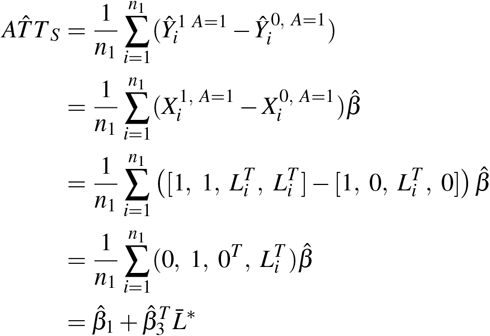

where 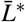 is a vector of length *J* with elements equal to the sample means among the treated of the *J* confounding variables.

Now let 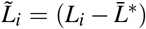 and 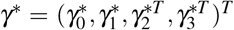, and consider finding 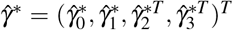 which solves

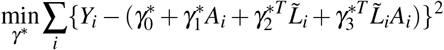

Rearranging this optimization problem similarly to A.3 and reparameterizing analogously to above yields 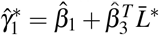. Note that 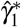 is the estimated exposure coefficient from the g-formula model proposed in the Supplementary Methods for the ATT, and the right side of this equality is exactly 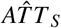 from above. Thus the equivalence of the two g-formula estimators has been shown for the ATT as well.

The proof for the equivalence of the average treatment effect in the untreated (ATU) is similar to the proof for the ATT.

## Appendix B R Workflows

## B.1 Generate Population Data and Analysis Dataset

This section contains R code for (i) generating a simulated dataset similar to those used in the simulation studies of the main text, and (ii) analyzing these data with each of regression, IPW, and the parametric g-formula to obtain the estimated effect of smoking in the smokers and the corresponding estimated standard errors. The dataset contains 770 individuals, which are sampled from a population of 10 million people.

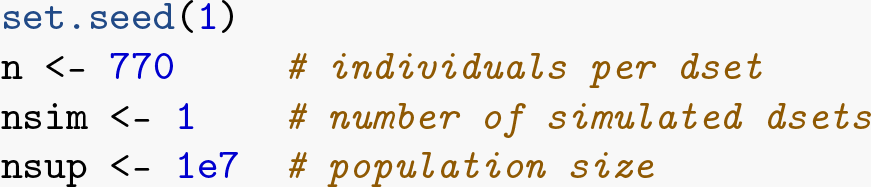

The following variables are generated for the population, according to the observed characteristics of the METSIM cohort.

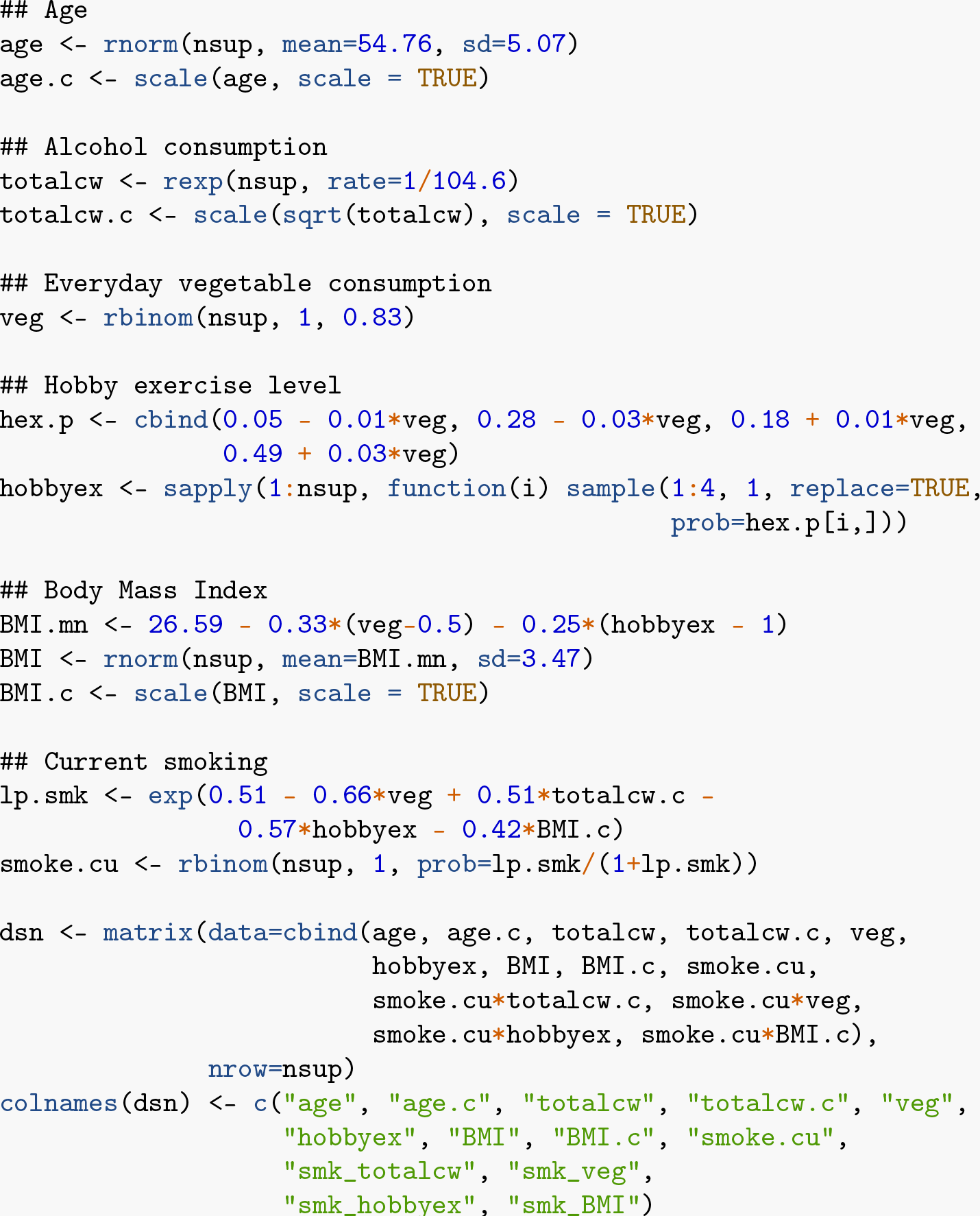

The dataset is assembled by sampling without replacement from the population. Note that if multiple datasets were being generated (i.e., `nsim`> 1), the same individual may be represented in more than one dataset. However, as written, this code doesn’t allow for individuals to be represented more than once within a single dataset.

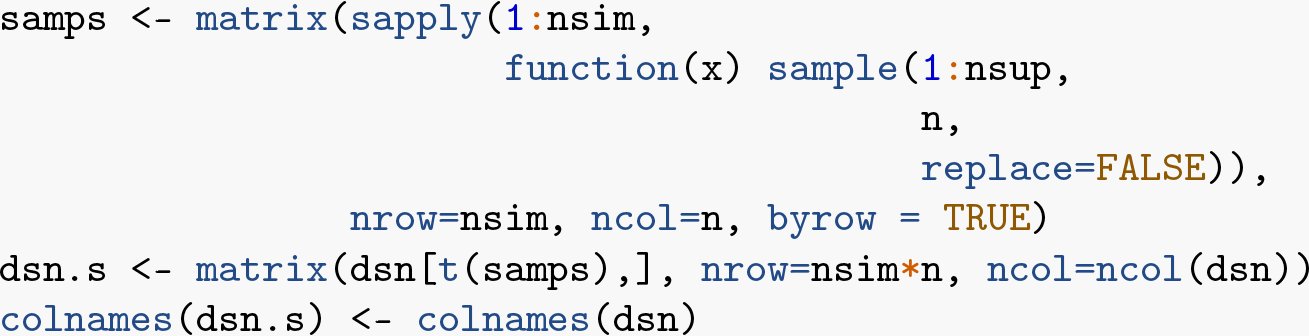

## B.2 Generate Gene Expression Data

Next, a matrix of coefficients is defined to set the influence of each of the covariates, the exposure, and their interactions on the average treatment effect in the treated. These six sets of coefficients correspond to the six simulation scenarios described in the main text.

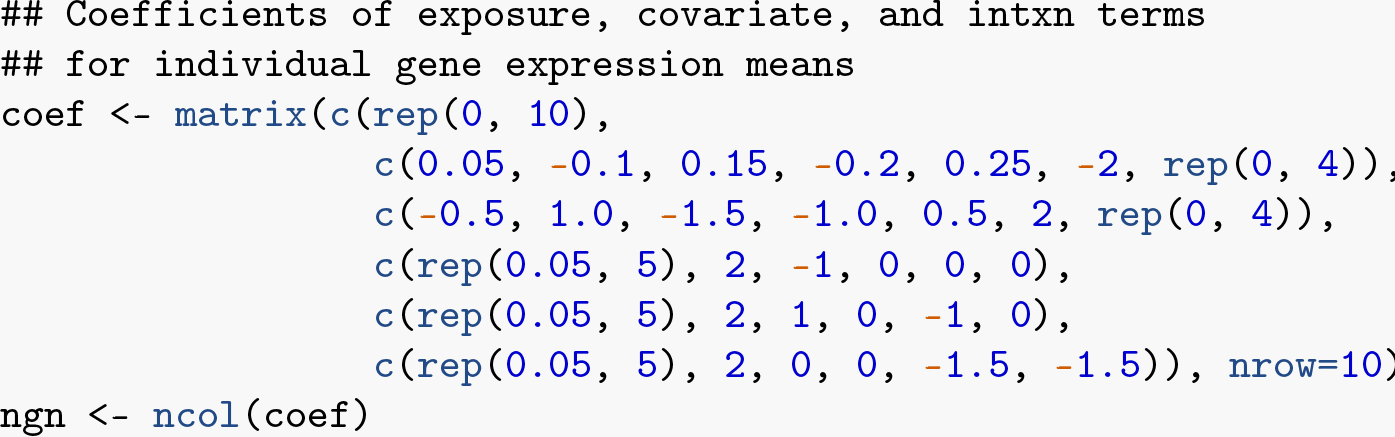

The individual covariate values are combined with the coefficients to produce a mean expression level for each gene in each individual. In other words, each person is given a mean expression level for each of the simulated genes based on their set of covariate and exposure values. These means are used to generate gene expression values from a normal distribution with standard deviation 0.24, for each gene.

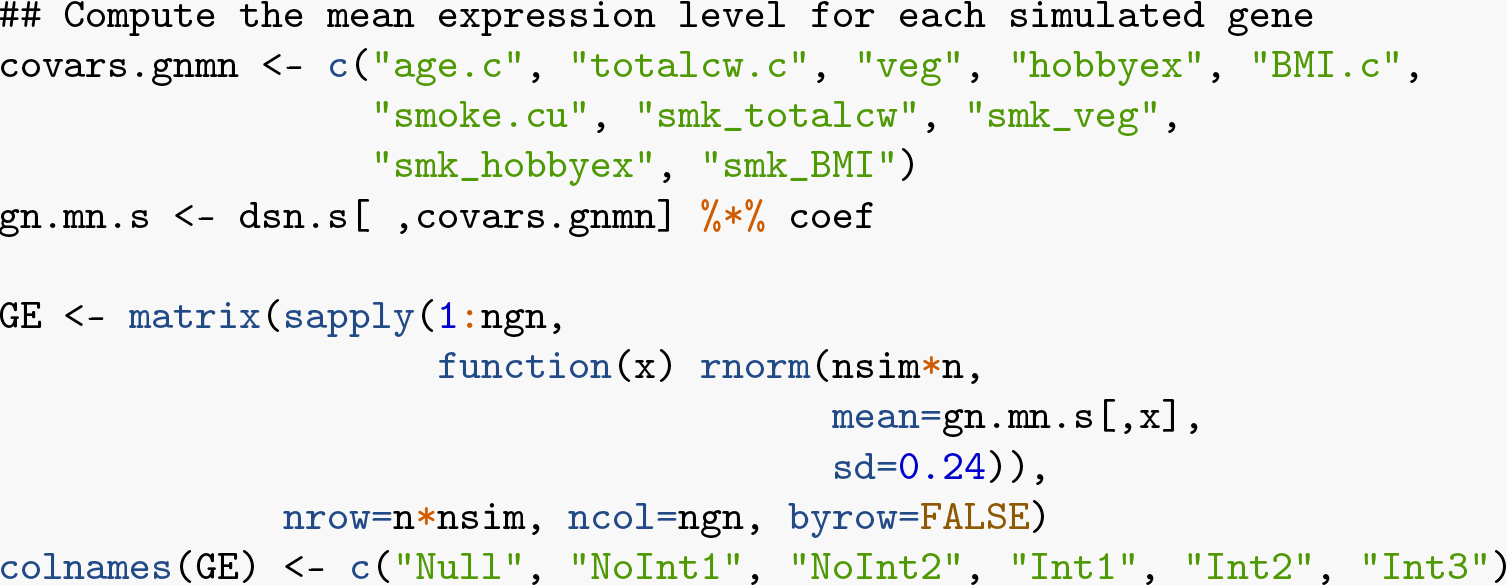

## B.3 Obtain True ATT for Each Gene

The true population ATT is computed for each gene by summing the contributions from each covariate. The covariate main effects don’t contribute to the ATT, only the exposure and the exposure-covariate interaction terms. Since the ATT is of interest, the contributions made by the interaction terms are the product of the coefficients and the mean value of each covariate in the current smokers (i.e., in the exposed). The contribution from the exposure is just the coefficients themselves.

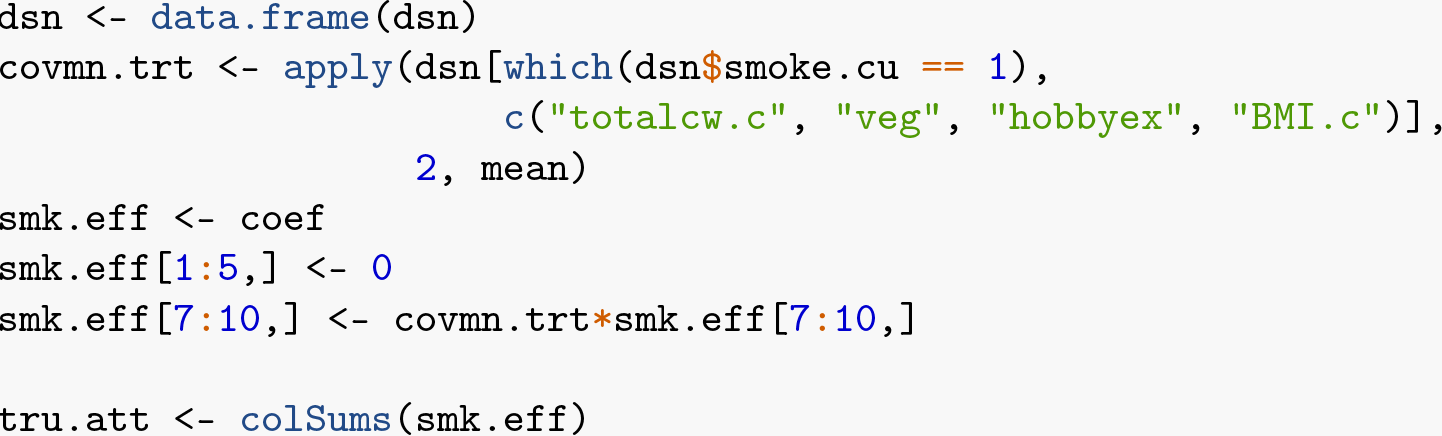

## B.4 Analyze Simulated Dataset

Analyses are conducted according to the models used in the main text for each method. This includes the addition of the quadratic term for age, though in simulations this term doesn’t contribute to the ATT. For each method, the estimates and their estimated standard errors can be used to construct Wald confidence intervals and obtain *t*-statistics and p-values.

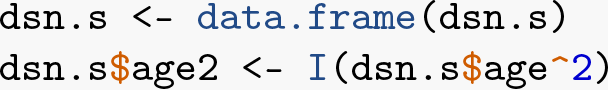

## B.4.1 Traditional Regression Analysis

The linear regression model is fit to the data once per gene using the `lmFit` function from the `limma` package for convenience and speed. Note that the standard errors recorded are the usual least squares estimates, not the moderated standard errors that the `lmFit` function also produces.

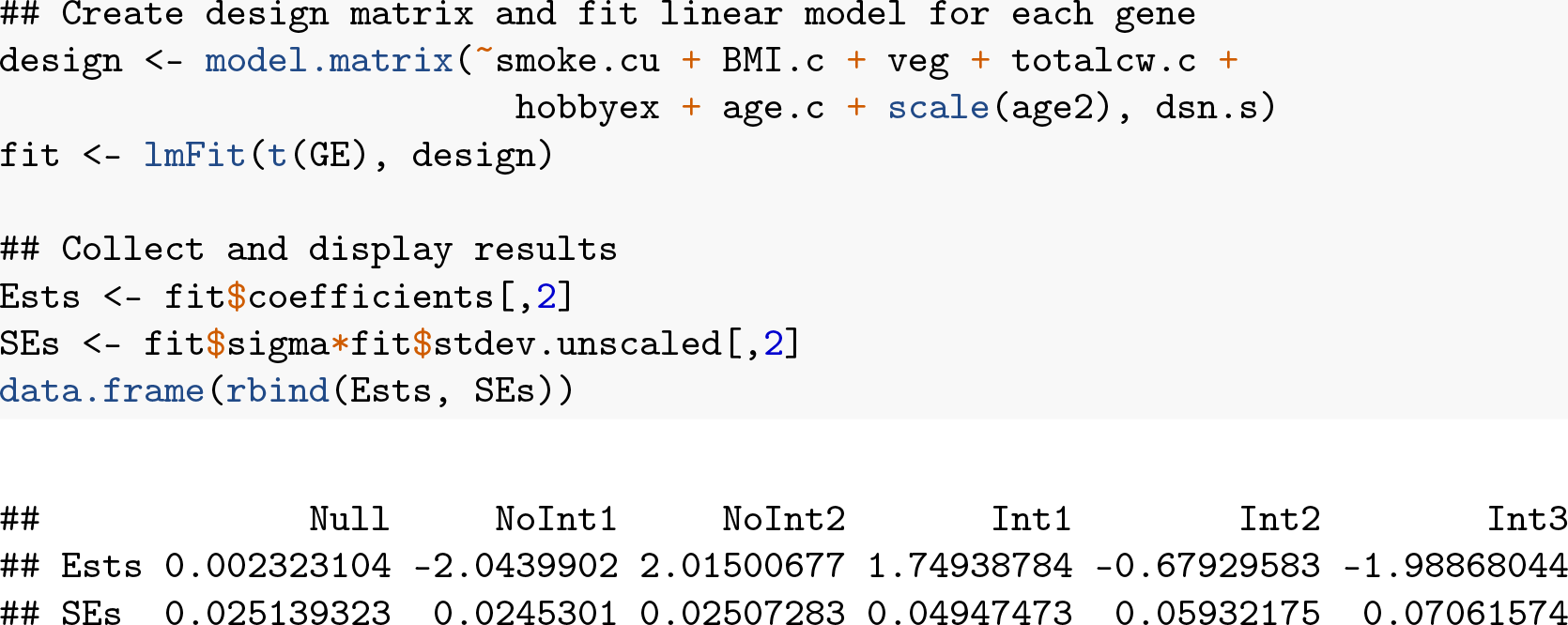

## B.4.2 Inverse Probability Weighting Analysis

Begin the IPW analysis by fitting a logistic regression model of the exposure to obtain parameter estimates needed for constructing the IP weights. Once the weights have been computed, it is good practice to check their distribution for extreme values and to ensure the mean is close to its expected value 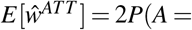 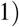. For this dataset, there appear to be no extreme values and the mean is very close to its expected value.

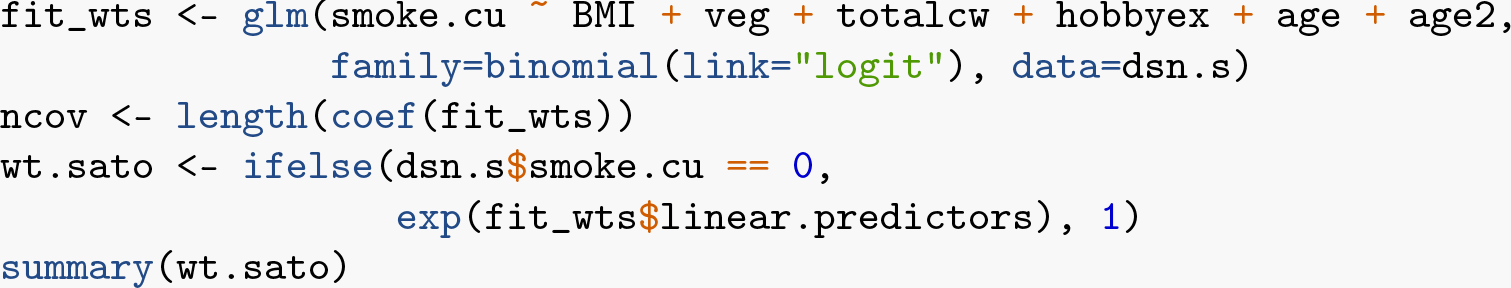
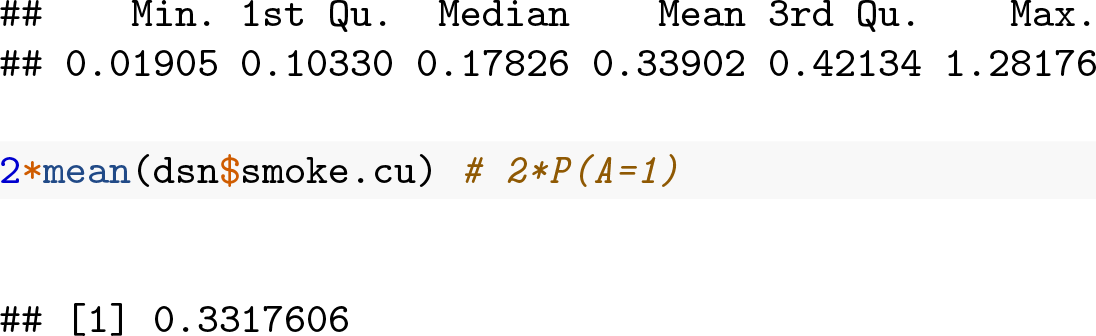

A simple linear regression model is fit for each gene with the weights, again using the `lmFit` function. The estimated ATT for each gene is recorded.

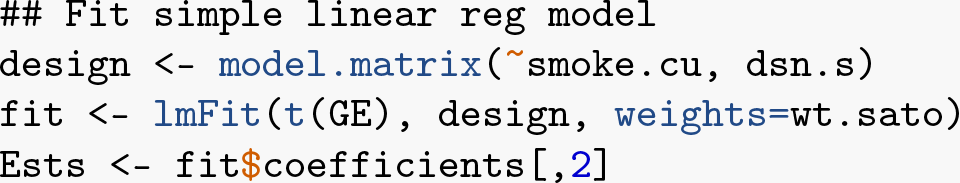

The standard error estimates from ordinary least squares are not appropriate in general for the IPW estimator, so the `geex` package is used to compute consistent standard error estimates. This approach stacks the estimating equations for the weights with those for the Hajek estimator using the function `estfun_IPW`, thereby accounting for estimation of the weights. The `m_estimate` function of the `geex` package computes the variance-covariance matrix for all estimated quantities, and the elements corresponding to the Hajek estimator are selected to compute the standard error estimate for each gene.

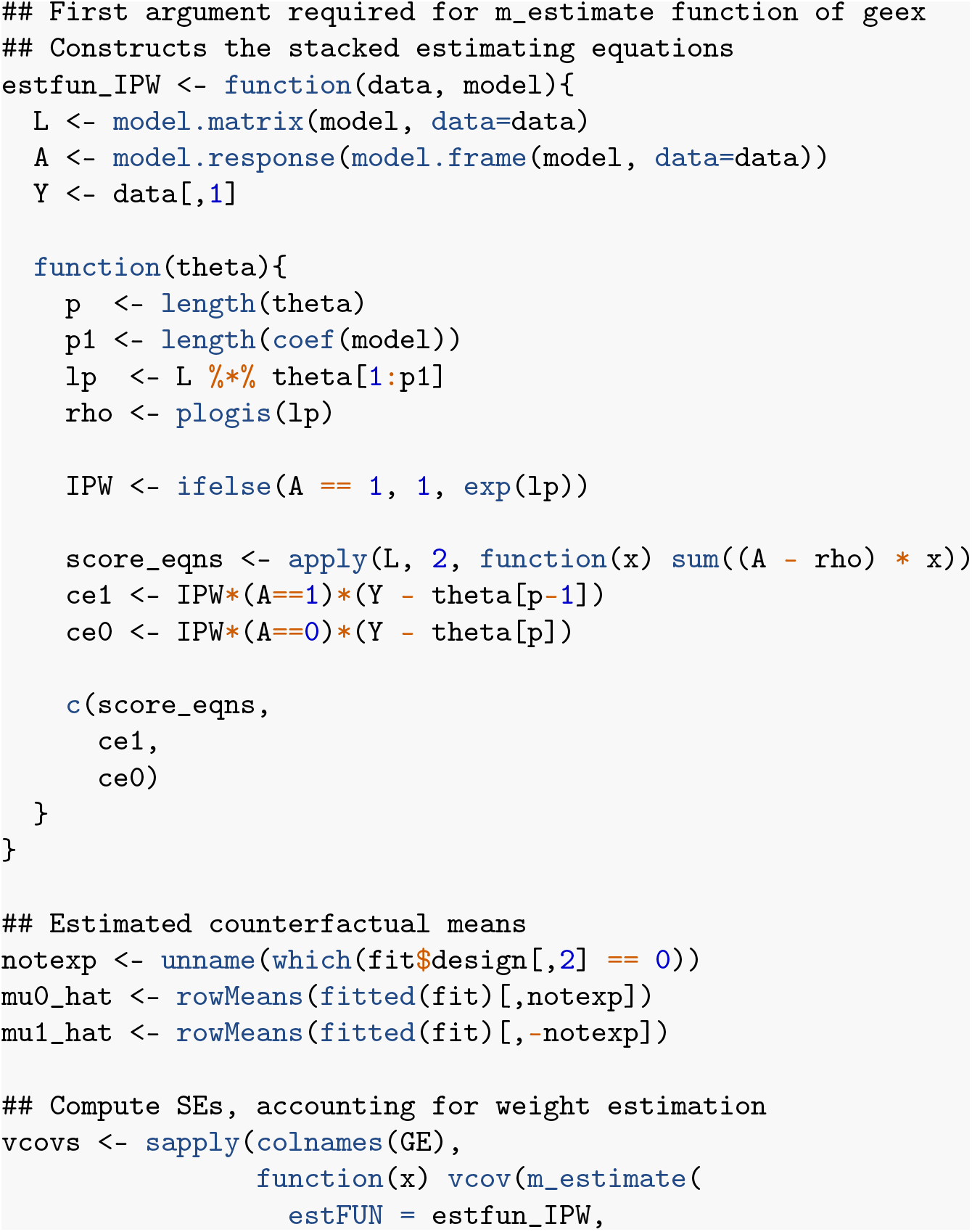
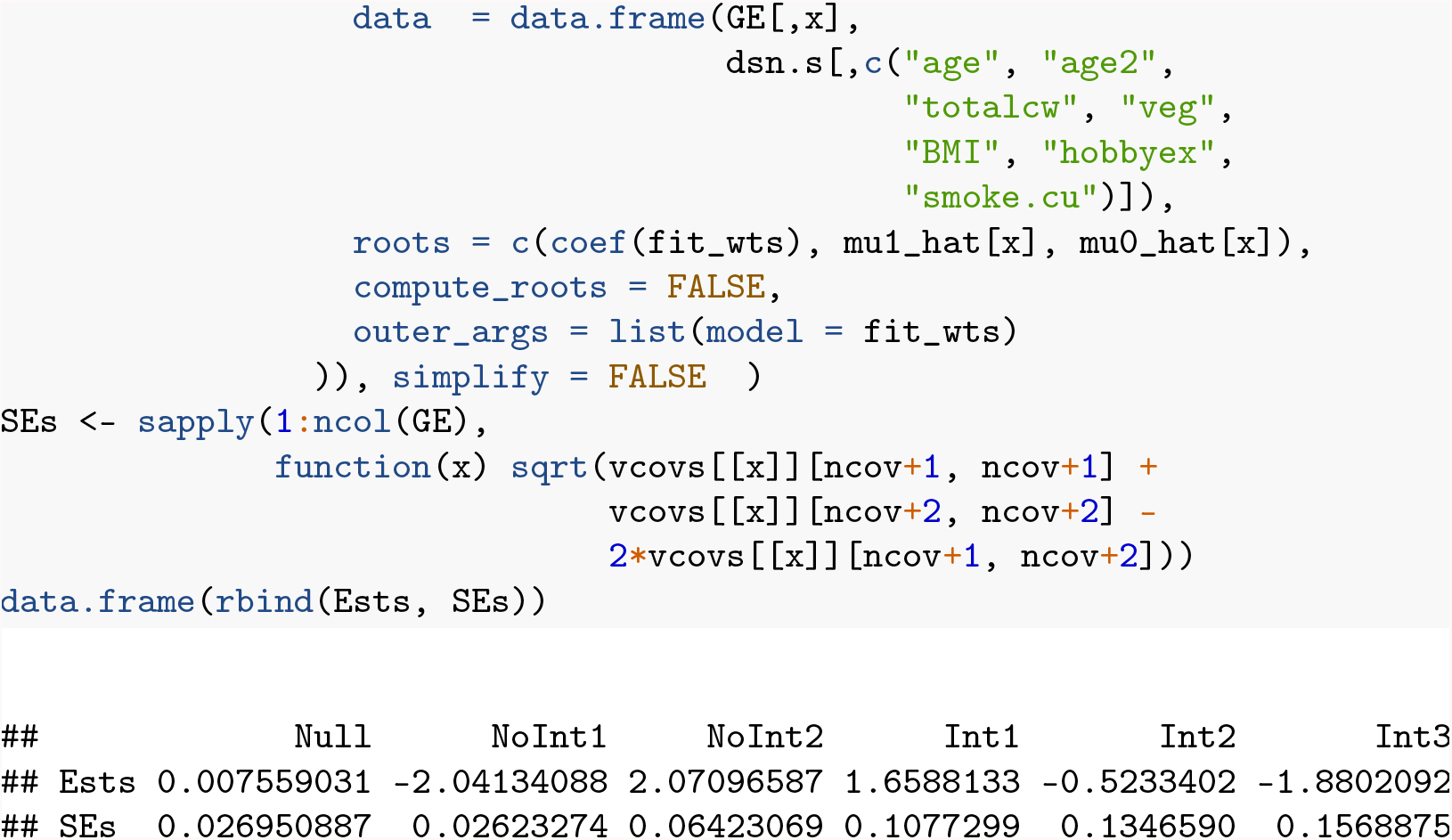

## B.4.3 Parametric G-formula Analysis

Fitting the g-formula model requires first centering all non-exposure covariates at their sample mean in the exposed. Once the appropriately centered main effects and interactions are generated, the full regression model is fit to obtain the estimated ATT for each gene, once more using `lmFit`. As with IPW, the standard error estimates are computed using stacked estimating equations to take the estimation of the covariate means into account.

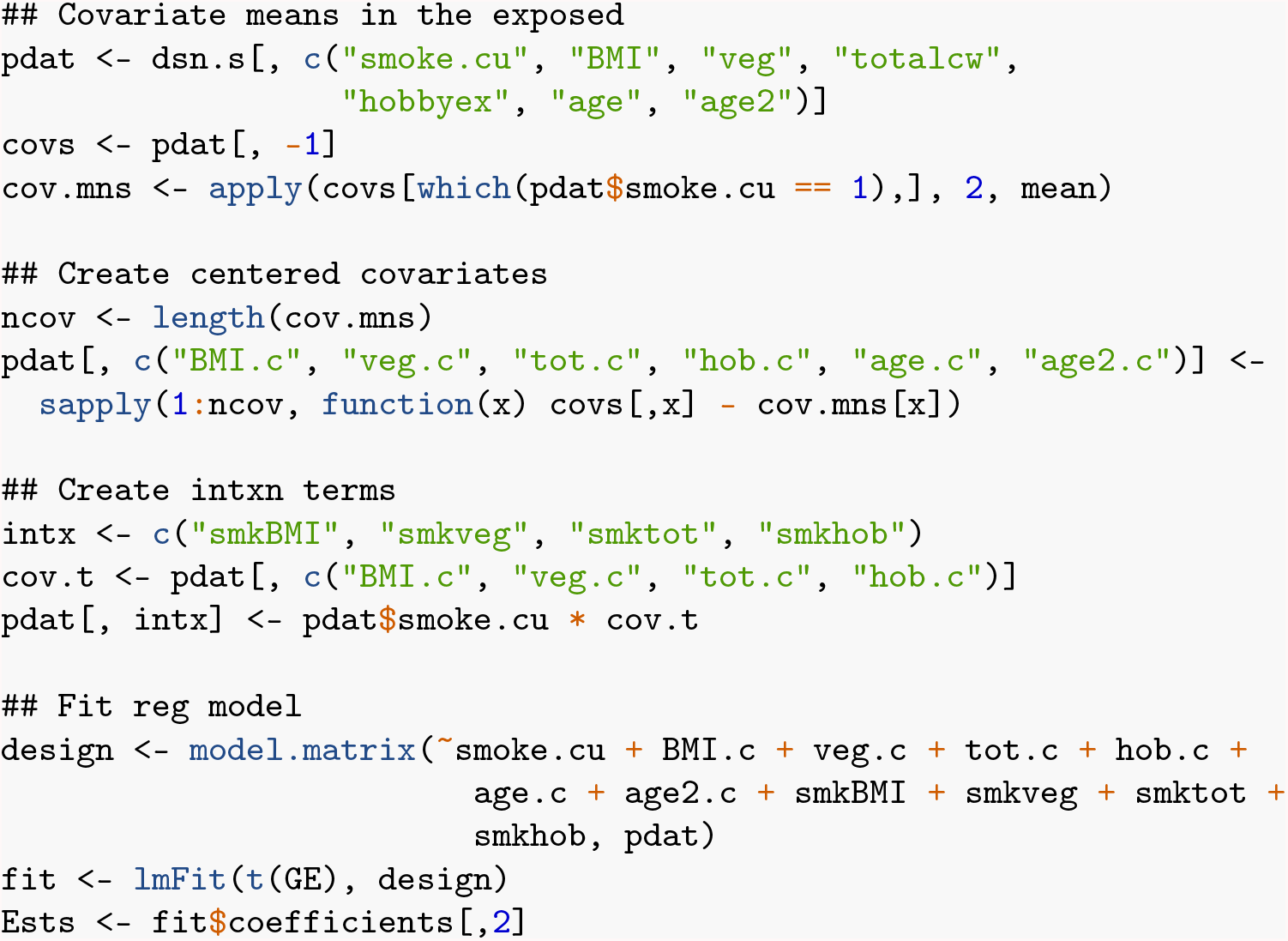

Similarly to above, the `estfun_gf` function is required by geex to produce the set of stacked estimating equations. The `m_estimate` code is analogous to that shown above for IPW, but here the standard error estimate of the g-formula estimator can be obtained directly from the diagonal of the variance-covariance matrix for each gene.

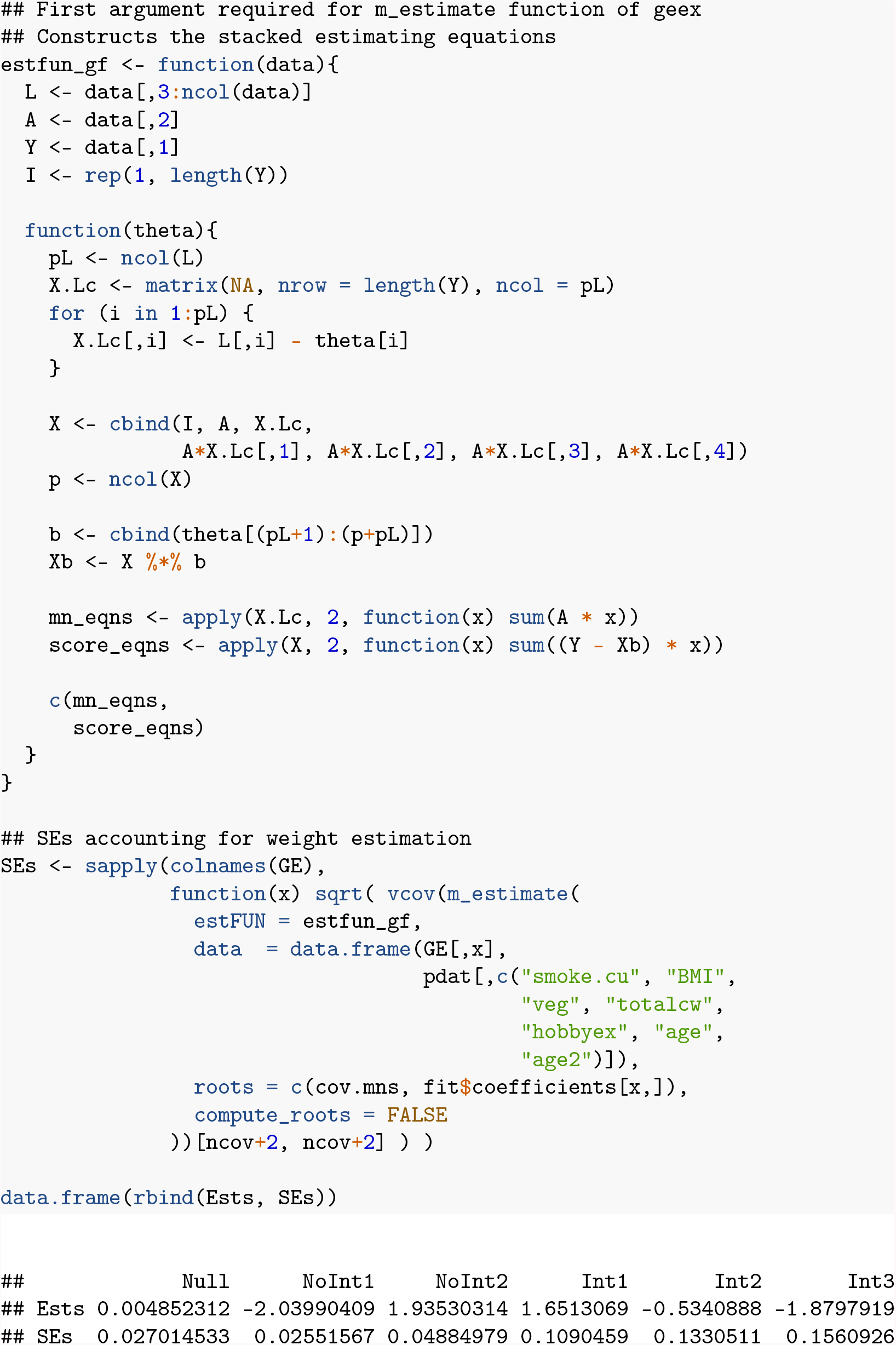

## Supplementary Methods

### Data Preprocessing

Microarray data (Affymetrix Human Genome U219 Array) from the METSIM project (Laakso et al., 2017) was downloaded from GEO accession GSE70353 (Civelek et al., 2017) using the Bioconductor package GEOquery (Davis and Meltzer, 2007). The downloaded data was normalized by the study authors. Microarray measurements per probeset were summarized for each gene using the median polish method from Tukey (1977) (the medpolish function in the R programming environment) on log_2_ transformed, normalized expression data.

In general, missing covariate values in the data set must be addressed before employing regression, IPW, or the parametric g-formula. If the percentage of individuals with missing covariates is low and the data are believed to be missing completely at random (MCAR), then a complete case analysis may be expected to not introduce bias. However, it is rarely the case that both of these criteria are met, and so it is recommended to take a more sophisticated approach such as multiple imputation (Moodie et al., 2008; Perkins et al., 2018). For further details and recommendations on handling missing covariates data when fitting causal models, see Moodie et al. (2008).

Assume for all of the following models that *Y*_*g*_ have been log_2_ transformed and normalized, and that all probe sets have been collapsed (e.g., using median polish) resulting in one measure per gene per subject. All vectors are assumed to be column vectors throughout.

### Comparing Approaches for Exposure Effect Estimation

#### Regression

For modeling gene expression *Y*_*g*_ of gene *g* as a function of some exposure of interest *A* in the presence of confounding, linear regression is the conventional approach. All potential confounders *L*, where *L* is a vector of length *J*, were included in the model as covariates along with the exposure variable *A*. In particular, the model can be written

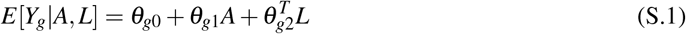

for each gene *g*, where *θ*_*g*2_ is also a vector of length *J*. The estimated exposure effect 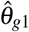 for gene *g* and its estimated standard error were computed in the usual fashion using ordinary least squares. Note that this model can be fit with or without interaction terms; here interactions were omitted from the model.

Ultimately the goal is to obtain consistent estimates and valid confidence intervals for the exposure effect in the exposed on gene expression, i.e., the ATT. The parameter estimate 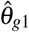 here is interpretable as the exposure effect in the population, not as the exposure effect in the exposed, unless operating under the assumption that the ATE and ATT are equal.

#### Inverse Probability Weighting

Inverse probability weights were computed by fitting the following logistic regression model with the binary exposure *A* as the outcome and the set of *J* potential confounders *L* as the predictors.

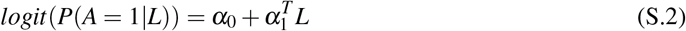

where *α*_1_ is a vector of length *J*. The fitted values from this logistic regression model were used to construct the individual weights that were then used in the models for gene expression. Choice of weights depends on the target population; this paper focuses on the ATT, so the weights given below first derived in Sato and Matsuyama (2003) were used. These weights take the form for each individual *i* of the ratio of the conditional probability of the subject being exposed to the conditional probability of the subject’s actual exposure status. That is, the weight for subject *i* equals

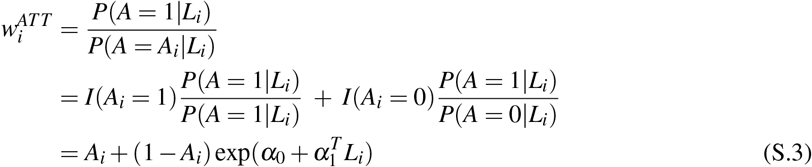

So, if a subject was exposed their weight was simply equal to one. For unexposed subjects, weights were estimated by substituting in the estimates 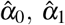 from fitting model S.2 for the true parameter values in S.3. When using IPW it is good practice to check that the mean of the weights is close to their expected value; for these weights, expect 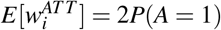. If interested in the ATE instead, see Robins et al. (2000).

After estimating the weights, the linear regression model

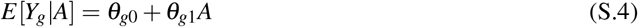

was fit for each gene *g* using weighted least squares with weights 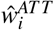, yielding

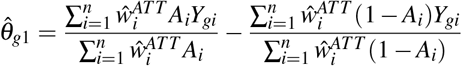

This estimator is sometimes referred to as the Hajek or modified Horwitz-Thompson estimator (Hernán and Robins, 2020), and is consistent for the ATT for each gene. Note that consistency of the Hajek estimator depends on the model for *A*|*L* in S.2 being correctly specified. No outcome model is assumed; fitting S.4 by weighted least squares is simply a convenient way to compute the Hajek estimator using standard software.

#### Parametric g-formula

In this final approach, the following altered version of the initial linear regression model was fit to the data.

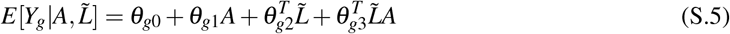

where 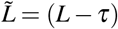 and *τ* is a vector of constants of length *J*, and *θ*_*g*2_, *θ*_*g*3_ are parameter vectors of length *J*. This model differs from model S.1 in two key ways: all first order interactions between *A* and the covariates *L* have been added, and all covariates *L* have been centered at *τ*. Since the ATT was of primary interest here, *τ* was chosen to contain the means of the covariates *L* among the exposed, *E*[*L*|*A* = 1]. Since the true means are unknown, a consistent estimate for *τ* was substituted, namely 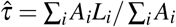. If interested in the ATE instead, let *τ* = *E*[*L*] and consistently estimate using 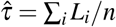.

The model was then fit using ordinary least squares to obtain the estimated ATT for gene *g*, 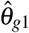. This estimator is equivalent to the estimator Snowden et al. (2010) demonstrated, which is computed in the *CAUSALTRT* procedure in SAS, and is thus consistent for the ATT; details in Appendix A.

### Standard Error Estimators for Each Method

The standard errors for the exposure effect estimator from the linear regression model were obtained using ordinary least squares, in keeping with the conventional approach. Estimating equations were used to compute the standard error of both the IPW Hajek estimator and the parametric g-formula estimator (Stefanski and Boos, 2002).

When taking an estimating equations approach to computing the standard errors, for IPW one must decide whether or not to take the estimation of the weights into account. The variance estimator that results from treating the weights as known is referred to here as the robust sandwich variance estimator (robust SVE). Computing the robust SVE is readily accomplished using various R packages such as *sandwich* or *geepack*, or using the *GENMOD* procedure in SAS. Accounting for the weight estimation in the variance computation, on the other hand, is the default if using the *CAUSALTRT* procedure in SAS or can be accomplished through supplying the set of estimating equations to the *geex* package (Saul and Hudgens, 2017) in R.

When using IPW and the ATE is of interest, there is a well-known result (Lunceford and Davidian, 2004) which states that the robust SVE is conservative when the weights are assumed known and consistent when weight estimation is taken into account. If using IPW to obtain the ATT, it is known from the theory of M-estimation (Stefanski and Boos, 2002) that this variance estimator is consistent when weight estimation is taken into account; however, it can be either conservative or anti-conservative when weights are assumed fixed. If computing the standard errors with *geex*, the set of estimating equations needed include the score equations from the logistic regression model in S.2 along with the two estimating equations corresponding to the two pieces of the Hajek estimator.

Bootstrapping is commonly employed to obtain standard errors when the g-formula is used to obtain the ATE or ATT (Efron and Tibshirani, 1986; Snowden et al., 2010; Wang et al., 2017), which is a valid option, but using stacked estimating equations provides a closed-form alternative. It is recommended when using the estimating equations approach that the covariate mean estimation be taken into account; again, by estimating equation theory, these standard errors are consistent and yield valid confidence intervals. If using *geex* to compute the standard errors for this estimator, the set of estimating equations needed are those corresponding to the estimation of each covariate mean and the parameters in model S.5.

In the simulations and data analyses of the paper, the variance of the ATT using the IPW estimator was calculated both ways and the two estimates were found to be fairly different in some instances and nearly identical in others. The same approach was taken for the variance of the ATT using the g-formula estimator, and the standard errors were substantially larger when accounting for estimation of covariate means. The standard error estimates reported were computed taking into account the estimation of the weights and the covariate means. The R markdown workflow that accompanies this paper includes code for computing the variance using stacked estimating equations for both IPW and the g-formula.

The standard errors for all methods were used to construct Wald 95% confidence intervals and perform *t*-tests of *H*_0_ : *θ*_*g*1_ = 0, i.e., no effect of exposure on gene expression in the exposed for gene *g*.

### Assumptions

With all methods presented here, it is assumed the gene expression data have already been normalized and reduced to one observation per person per gene (i.e., not probe-level data). These methods also require that there are no missing values; if missing values are present in the covariates *L* or exposure *A*, see the recommendations in the section above. For these methods to yield consistent estimates, it is also assumed that there is no bias due to selection or measurement error. Importantly, formal arguments for IPW and the parametric g-formula involve asymptotic justifications, and there is no guarantee that these methods will perform well for small or moderate sample sizes (e.g., *n* < 40 for IPW (Pirracchio et al., 2012)).

In order for the IPW and g-formula methods to adequately adjust for confounding, the set of covariates *L* must satisfy the conditional exchangeability assumption (*Y*^*a*^ ⊥ *A*|*L*). It is also required that positivity *P*(*A* = *a*|*L* = *l*) > 0 for all *l* where *dF*_*L*_(*l*) > 0 and *F*_*L*_ is the CDF of *L*, and the Stable Unit Treatment Value Assumption (SUTVA) hold. SUTVA requires causal consistency, i.e., no different versions of exposure, and no interference, i.e., one individual’s exposure status doesn’t affect another individual’s gene expression.

For the IPW and g-formula estimators to yield consistent effect estimates, the above assumptions must hold. When using IPW, the additional assumption that the model of *A*|*L* is correctly specified is needed as well. For the g-formula, no specification of a model for *A*|*L* is needed, but the model for *Y*|*A, L* must be correctly specified.

## Supplementary Results

### Simulation Study for Ordinary vs Moderated *t* Statistic

In observational genomics studies, the total sample size is often large enough for results depending on large sample theory to hold. The *limma* package (Smyth, 2004) is used in standard practice to obtain effect estimates, *t*-statistics, and p-values for each gene when assessing the effect of some exposure on gene expression. This package computes a moderated *t*-statistic that is shown to perform well in small sample sizes, as are found in traditional genetic studies, and which converges to the ordinary *t*-statistic as the sample size increases. The moderation of this *t*-statistic comes into play with empirical Bayes moderation of the standard errors toward a common value; here it is shown empirically that these moderated standard errors are practically equivalent to the ordinary standard errors in large samples.

In Figure S.1 the difference between the moderated and ordinary *t*-statistics for an example gene are given for a variety of sample sizes, and with balanced and unbalanced group assignment. In particular, a randomly chosen gene from the METSIM cohort data was used, and the 770 participants were randomly sampled according to their current smoking status to create the analysis datasets. For each combination of sample size and group allocation, 200 analysis datasets were constructed. Samples were drawn such that the same individual may have been represented in more than one dataset, but not more than once within a single dataset. Both types of *t*-statistic and their difference were computed for each dataset, and boxplots of the 200 differences are given for each scenario in Figure S.1. Regardless of the group allocation, once the total sample size was around 100 or larger, the moderated and ordinary *t*-statistics were practically equivalent.

**Figure S1:**
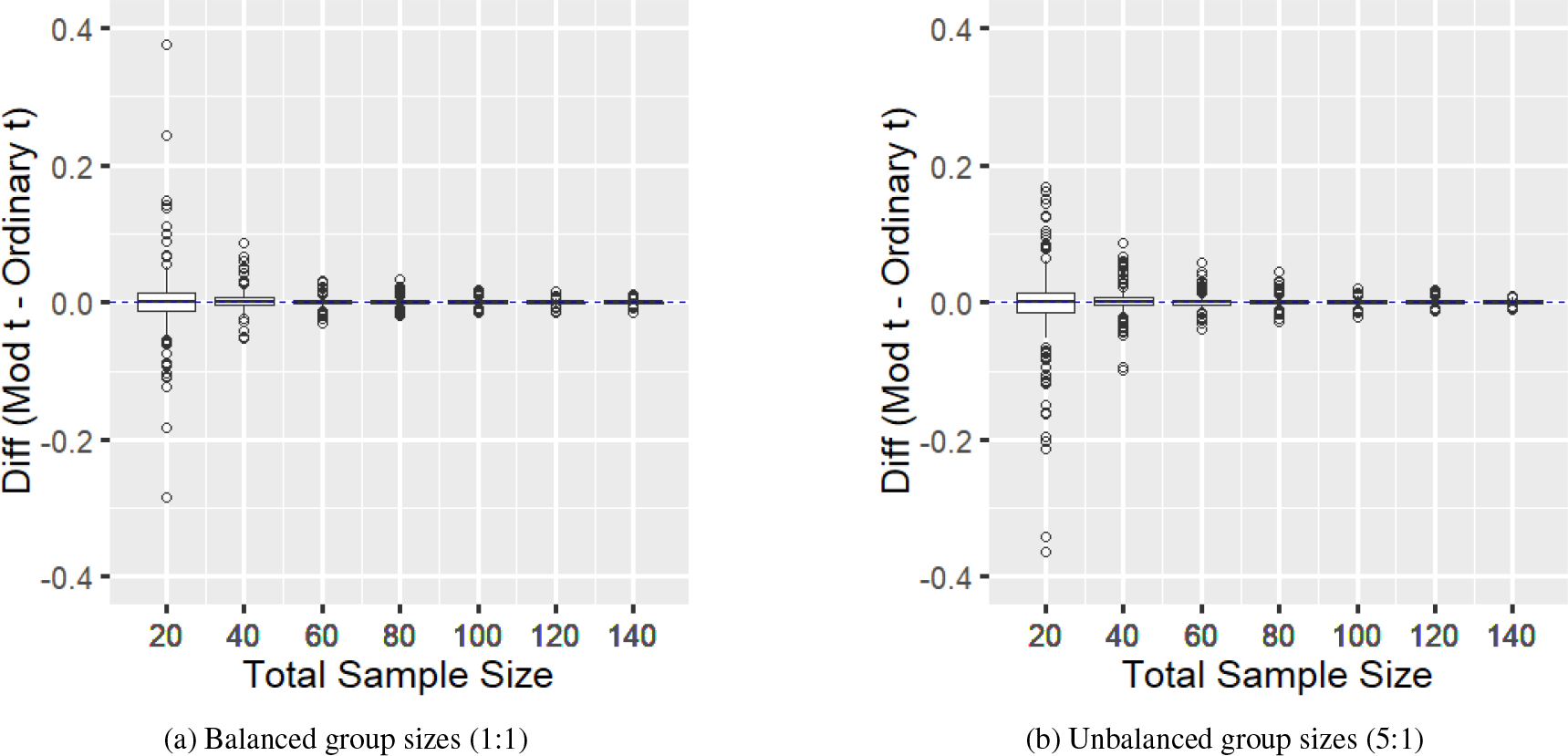
Difference between the moderated and ordinary *t*-statistics for various sample sizes with (a) balanced (1 non-smoker : 1 smoker) and (b) unbalanced groups (5 non-smokers : 1 smoker). The boxplot for each sample size represented in (a) and (b) summarizes the difference in *t*-statistics for 200 datasets, each derived from the same gene.

All analyses in this paper were conducted with well over 100 individuals, and so ordinary linear regression was used for simplicity.

### Sensitivity Analyses of METSIM Microarray Data

As mentioned in the main text, there was one large IP weight from the METSIM primary analysis whose influence deserves further investigation. Deleting this individual from the data resulted in a sample of size 769, which was analyzed again in the same manner as above. Results of the regression, IPW, and g-formula sensitivity analysis are compared to the results from the primary analysis in Figure S.2a - S.2c.

Additionally, the leverage values for each individual were computed from the design matrices of the g-formula and regression methods. The individual with highest leverage value was the same for both methods, and another sensitivity analysis was performed by deleting this individual; the results of this second sensitivity analysis are compared to the results from the primary analysis in Figure S.2d - S.2f. Notably, the individual with second highest leverage value was the same individual who generated the largest weight and who was deleted in the sensitivity analysis above. The genes represented in this figure are the same set as those in Figure 1, namely the top 50 genes as ranked in the primary analysis.

**Figure S2:**
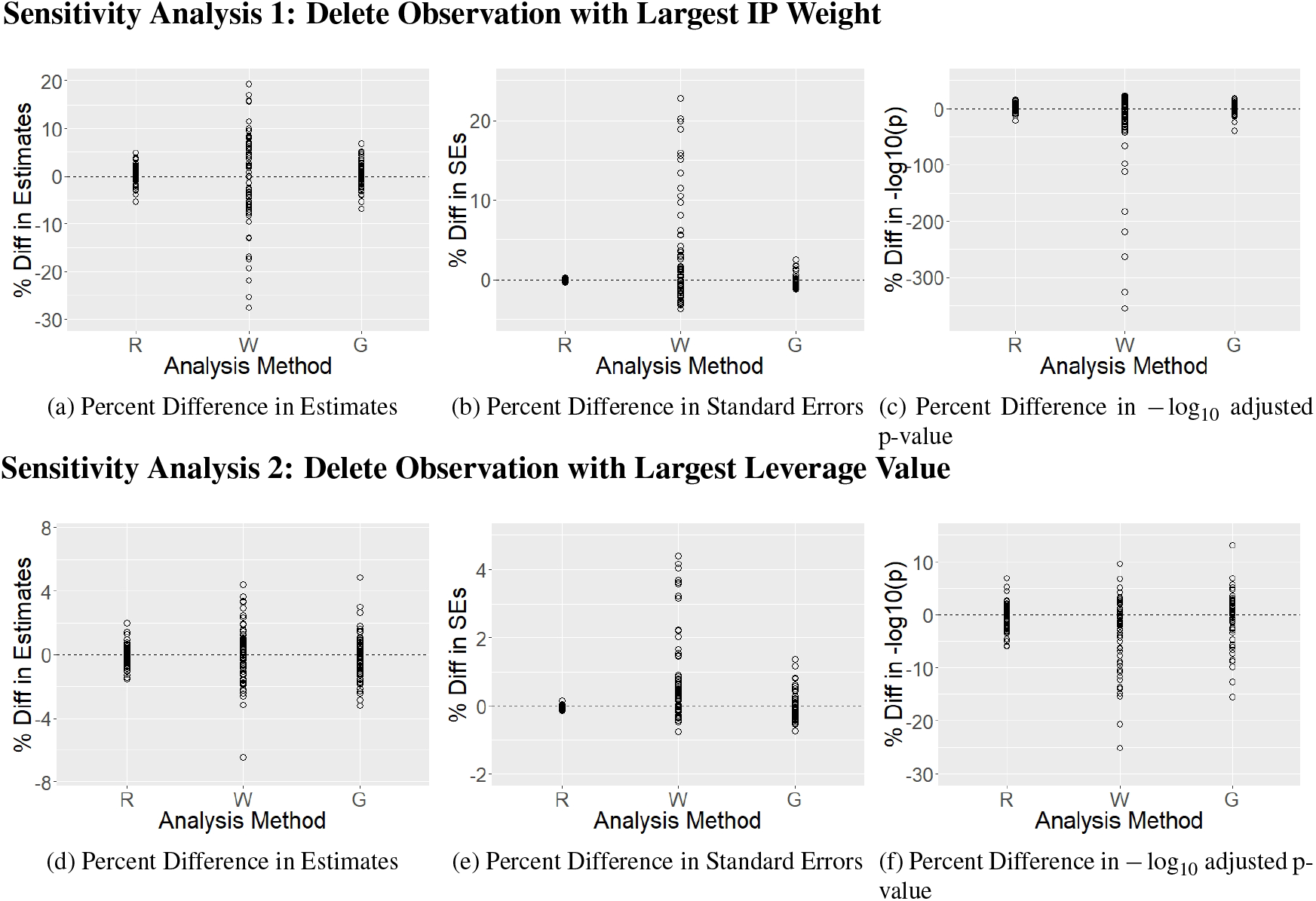
(a)-(c): Comparison of METSIM primary and sensitivity analysis results when deleting observation with largest weight. Top 50 genes are represented, ranked by p-value. (a), (b), and (c) respectively show the percent difference (primary - sensitivity) of the effect estimates, standard errors, and −log_10_ Benjamini-Hochberg adjusted p-values. (d)-(f): Comparison of METSIM primary and sensitivity analysis results when deleting observation with largest leverage value, which was the same observation for both the regression and g-formula methods. Top 50 genes are represented, ranked by p-value. (d), (e), and (f) respectively show the percent difference (primary - sensitivity) of the effect estimates, standard errors, and −log_10_ Benjamini-Hochberg adjusted p-values. R = Regression, W = Inverse Probability Weighting, G = Parametric G-Formula.

In particular, the first row of this figure shows the deletion of the observation with largest weight had very little effect on the regression and g-formula estimates, standard errors, and p-values. On the other hand IPW appears to be more sensitive to the deletion of this observation, with percent difference in effect estimates and standard errors ranging up to a magnitude of 30 and 25 respectively. These changes were reflected in the −log_10_ adjusted p-values as well; the bulk of the IPW p-values did not change by more than a magnitude of 50 percent, but p-values for some estimates changed by more than a magnitude of 300 percent. While the effect on the majority of the IPW estimates, standard errors, and p-values was small, some of the top 50 genes saw substantial changes.

For the second sensitivity analysis where the observation with largest leverage value was deleted, the g-formula and IPW estimates were affected similarly ((−8, 6) percent difference) and to a slightly larger degree than the regression estimates ((−2, 3) percent difference). The change in standard errors was largest again for IPW but still contained to (−1, 5) percent difference, much smaller than for the previous sensitivity analysis. The change in g-formula standard errors was contained to (−1, 2) percent difference, and the regression standard errors changed by less than one percent in either direction. These changes were reflected in the −log_10_ adjusted p-values as well, with the percent difference for the regression and g-formula p-values having a spread comparable to the previous sensitivity analysis, and with the IPW p-values being considerably less variable than before.

## Notes

**Funding Information:** S.A.R. was supported by The Chancellor’s Fellowship from The Graduate School at the University of North Carolina at Chapel Hill. M.G.H. was supported by R01 AI085073. M.C. was supported by R01 DK118287 and R21 HL135230. K.L.M. was supported by R01 DK093757. M.I.L. was supported by R01 HG009937, R01 MH118349, and P30 ES010126.

## References

Anders, S. and Huber, W. (2010). Differential expression analysis for sequence count data. Genome Biology, 11(10):R106.

Benjamini, Y. and Hochberg, Y. (1995). Controlling the False Discovery Rate : A Practical and Powerful Approach to Multiple Testing. Journal of the Royal Statistical Society. Series B (Methodological), 57(1):289–300.

Civelek, M., Wu, Y., Pan, C., Raulerson, C. K., Ko, A., He, A., Tilford, C., Saleem, N. K., Stančáková, A., Scott, L. J., Fuchsberger, C., Stringham, H. M., Jackson, A. U., Narisu, N., Chines, P. S., Small, K. S., Kuusisto, J., Parks, B. W., Pajukanta, P., Kirchgessner, T., Collins, F. S., Gargalovic, P. S., Boehnke, M., Laakso, M., Mohlke, K. L., and Lusis, A. J. (2017). Genetic Regulation of Adipose Gene Expression and Cardio-Metabolic Traits. American Journal of Human Genetics, 100(3):428–443.

Cui, S., Ji, T., Li, J., Cheng, J., and Qiu, J. (2016). What if we ignore the random effects when analyzing RNA-seq data in a multifactor experiment. Statistical Applications in Genetics and Molecular Biology, 15(2):87–105.

Davis, S. and Meltzer, P. S. (2007). GEOquery: a bridge between the Gene Expression Omnibus (GEO) and BioConductor. Bioinformatics, 23(14):1846–1847.

Efron, B. and Tibshirani, R. (1986). Bootstrap Methods for Standard Errors, Confidence Intervals, and Other Measures of Statistical Accuracy. Statistical Science, 1(1):54–75.

Gagnon-Bartsch, J. A. and Speed, T. P. (2012). Using control genes to correct for unwanted variation in microarray data. Biostatistics, 13(3):539–552.

Hejazi, N., Phillips, R., Hubbard, A., and van der Laan, M. (2018). methyvim: Targeted, robust, and model-free differential methylation analysis in r [version 1; peer review: 1 approved with reservations, 2 not approved]. F1000Research, 7(1424).

Hejazi, N. S., Kherad-Pajouh, S., van der Laan, M. J., and Hubbard, A. E. (2017). Variance stabilization of targeted estimators of causal parameters in high-dimensional settings. arXiv e-prints, page arXiv:1710.05451.

Heller, R., Manduchi, E., and Small, D. (2008). Matching Methods for Observational Microarray Studies. Bioinformatics, 25(7):904–909.

Hernán, M. and Robins, J. (2020). Causal Inference. Chapman & Hall/CRC, Boca Raton.

Laakso, M., Kuusisto, J., Stančáková, A., Kuulasmaa, T., Pajukanta, P., Lusis, A. J., Collins, F. S., Mohlke, K. L., and Boehnke, M. (2017). The Metabolic Syndrome in Men study: A resource for studies of metabolic & cardiovascular diseases. Journal of Lipid Research, 58(3):481–493.

Law, C. W., Chen, Y., Shi, W., and Smyth, G. K. (2014). voom: precision weights unlock linear model analysis tools for rna-seq read counts. Genome Biology, 15(2):R29.

Leek, J. T. and Storey, J. D. (2007). Capturing heterogeneity in gene expression studies by surrogate variable analysis. PLOS Genetics, 3(9):1–12.

Lunceford, J. K. and Davidian, M. (2004). Stratification and weighting via the propensity score in estimation of causal treatment effects: A comparative study. Statistics in Medicine, 23(19):2937–2960.

Moodie, E. E., Delaney, J. A., Lefebvre, G., and Platt, R. W. (2008). Missing confounding data in marginal structural models: A comparison of inverse probability weighting and multiple imputation. International Journal of Biostatistics, 4(1):Article 13.

Moodie, E. E. M., Saarela, O., and Stephens, D. A. (2018). A doubly robust weighting estimator of the average treatment effect on the treated. Stat, 7(1):e205.

Naimi, A. I., Cole, S. R., and Kennedy, E. H. (2017). An introduction to g methods. International Journal of Epidemiology, 46(2):756–762.

Naimi, A. I. and Kennedy, E. H. (2017). Nonparametric Double Robustness. arXiv e-prints, page arXiv:1711.07137.

Nebert, D. W. and Russell, D. W. (2002). Clinical importance of the cytochromes P450. Lancet, 360(9340):1155–1162.

Perkins, N. J., Cole, S. R., Harel, O., Tchetgen Tchetgen, E. J., Sun, B., Mitchell, E. M., and Schisterman, E. F. (2018). Principled Approaches to Missing Data in Epidemiologic Studies. American Journal of Epidemiology, 187(3):568–575.

Pirracchio, R., Resche-Rigon, M., and Chevret, S. (2012). Evaluation of the Propensity score methods for estimating marginal odds ratios in case of small sample size. BMC Medical Research Methodology, 12(70).

Robins, J. M., Hernán, M. A., and Brumback, B. (2000). Marginal structural models and causal inference in epidemiology. Epidemiology, 11(5):550–560.

Sato, T. and Matsuyama, Y. (2003). Marginal structural models as a tool for standardization. Epidemiology, 14(6):680–686.

Saul, B. C. and Hudgens, M. G. (2017). The Calculus of M-estimation in R with geex. arXiv e-prints, page arXiv:1709.01413.

Smyth, G. K. (2004). Linear Models and Empirical Bayes Methods for Assessing Differential Expression in Microarray Experiments. Statistical Applications in Genetics and Molecular Biology, 3(1):1–26.

Smyth, G. K., Michaud, J., and Scott, H. S. (2005). Use of within-array replicate spots for assessing differential expression in microarray experiments. Bioinformatics, 21(9):2067–2075.

Snowden, J. M., Rose, S., and Mortimer, K. M. (2010). Implementation of G-computation on a simulated data set: Demonstration of a causal inference technique. American Journal of Epidemiology, 173(7):731–738.

Stefanski, L. and Boos, D. (2002). The Calculus of M-Estimation. The American Statistician, 56(1):29–38.

Stegle, O., Parts, L., Piipari, M., Winn, J., and Durbin, R. (2012). Using probabilistic estimation of expression residuals (peer) to obtain increased power and interpretability of gene expression analyses. Nature Protocols, 7(3):500–507.

Sun, S., Hood, M., Scott, L., Peng, Q., Mukherjee, S., Tung, J., and Zhou, X. (2017). Differential expression analysis for RNAseq using Poisson mixed models. Nucleic Acids Research, 45(11):1–15.

Tukey, J. W. (1977). Exploratory Data Analysis. Addison-Wesley, Reading, PA.

Van De Wiel, M. A., Leday, G. G., Pardo, L., Rue, H., Van Der Vaart, A. W., and Van Wieringen, W. N. (2012). Bayesian analysis of RNA sequencing data by estimating multiple shrinkage priors. Biostatistics, 14(1):113–128.

Wang, A., Nianogo, R. A., and Arah, O. A. (2017). G-computation of average treatment effects on the treated and the untreated. BMC Medical Research Methodology, 17(1):1–5.

